# Environmental and spatiotemporal drivers of marine microbial communities from Antarctic and Subantarctic water masses

**DOI:** 10.64898/2026.08.13.742230

**Authors:** Manuel Ochoa-Sánchez, Jorge Acevedo, Yoshihiro Fujise, Tatsuya Isoda, Aída I. Murillo-Herrera, Eliana P. Acuña Gómez, Pedro Valenzuela, Claudio Moraga, Luis A. Pastene

## Abstract

The Southern Ocean harbors diverse marine microbial communities shaped by both local oceanographic conditions and dispersal limitations. However, this knowledge is mainly based on coastal Antarctic sites, whereas circumpolar Antarctic open sea and subantarctic ecosystems remain poorly explored. Here, we characterize marine microbial communities (using 16S rDNA high-throughput sequencing) and marine oceanographic data across two regions: the Subantarctic, involving two localities (the Magellan Strait and the Beagle Channel), and Antarctic open sea, involving two localities (Eastern Indian and Central South Pacific). We found extensive differences across regions and localities, characterized by distinct taxonomic patterns, alpha diversity, microbial composition, and enriched taxa profiles. Despite these differences, *Clade Ia*, *Amylibacter*, *NS5 marine group*, and *NS2b marine group* exhibited high prevalence across regions. Oceanographic parameters had variable relationships with microbial alpha diversity across regions: Sea surface temperature and salinity had a negative and positive correlation, respectively, in the Magellan Strait during 2024. In the Antarctic region, dissolved oxygen displayed a negative correlation in the Indian Ocean during 2024, whereas salinity displayed a more variable relationship in the Indian Ocean: positively correlated during 2024, while negatively correlated during 2025. Collectively, our results highlight a strong microbiological biogeographic structure in the Southern Ocean, both across broad scales (between Subantarctic and Antarctic regions) and within regions. Furthermore, our results show dynamic relationships between oceanographic variables and marine microbial diversity across Antarctic and Subantarctic regions.

## Introduction

The Southern Ocean (SO) is an enormous reservoir of marine biodiversity, encompassing waters below 40°S globally (Chapman et al., 2020). The SO displays multiple oceanographic features that differentiate it from northern polar marine ecosystems, such as iron scarcity in surface waters, distinct thermohaline circulation patterns due to the Antarctic Circumpolar Current (ACC), and lack of continental barriers (Boyd et al., 2010). Additionally, the seasonal input of freshwater and nutrients from sea ice and glacier melt is an important driver of marine productivity that promotes local hotspots of marine biodiversity (Boyd et al., 2000). These hotspots of marine productivity sustain complex trophic chains that harbor unique portions of global biodiversity (Cao et al., 2020; Griffith et al., 2026).

The diversity and composition of marine microbial communities from SO are heavily influenced by seasonal shifts (Iida et al., 2014; Liu et al., 2020). These seasonal microbial shifts shape the marine ecosystem dynamics, as they are linked to changes in dimethyl sulfoniopropionate (DMSP), which modulates sea surface temperature (O’Brien et al., 2022) as well as marine productivity and trophic dynamics (Arteaga et al., 2020; Christaki et al., 2021). Also, several studies have identified regionally specific microbial assemblages, highlighting the importance of biogeographic factors (Liu et al., 2020; O’Brien et al., 2022; Liu et al., 2024). These biogeographic patterns in marine microbial communities are likely structured by oceanographic fronts, which determine microbial diversity and composition across the SO. Nevertheless, evidence also supports microbial dispersal, which could be facilitated by marine currents (Schwob et al., 2021). Despite these insights, given the vast area that the SO encompasses, much of its associated microbial diversity remains largely unexplored.

In turn, subantarctic marine microbial diversity remains largely unexplored, particularly in southern South America (Ochoa-Sánchez et al., 2023a). Recent surveys of marine hosts that included marine samples have revealed spatial and temporal variability in microbial taxonomic patterns along the Magellan Strait. For example, in the eastern sector of the Magellan Strait, *Flavobacteraceae* and *Moraxellaceae* alternated in numerical dominance, whereas in southern areas of the Magellan Strait, *Flavobacteraceae* was the most abundant taxon despite interannual variability (Ochoa-Sánchez et al., 2024b, 2025a). Also, in two fjords of the Beagle Channel, Proteobacteria was the dominant fraction of the marine microbial community; however, freshwater discharges and oceanic influence were also important factors shaping microbial structure (Maturana-Martínez et al., 2021). However, most existing studies have focused on host-associated microbiotas or on specific fjords of South America; consequently, comprehensive investigations that provide a robust perspective on the subantarctic marine microbial diversity are still lacking.

The Magellan region is a subantarctic environment at the southernmost tip of South America, where the most iconic geological features are the Magellan Strait and the Beagle Channel, which are corridors connecting the Atlantic and Pacific Oceans. Both corridors originated during the Pleistocene glacial expansion, which created an intricate network of fjords and complex hydrographic patterns (De Muro et al., 2015; Ferreyra & González, 2024; Suárez et al., 2024). From the biological perspective, the Magellan Strait is a hotspot of biodiversity, like the Marine and Coastal Protected Area Francisco Coloane (CEQUA 2007), including several marine predators (Gibbons et al., 2000; Venegas et al., 2002; Ochoa-Sánchez et al., 2024a), while the Beagle Channel stands out as a unique glaciomarine system characterized by strong freshwater inputs from surrounding ice fields and glaciers, creating sharp environmental gradients that profoundly shape local marine communities (Rabassa et al. 2000).

The Antarctic open sea region, south of 60°S, considered in this study, is in the approximate position of the Antarctic Convergence (AC). Prydz Bay is located at the western boundary of this region, while the Ross Sea is located at its eastern boundary. This region is strongly influenced by the southern boundary of the ACC, which interacts with the coastal East Wind Drift (EWD) in a series of fronts and eddies. A series of gyres link the EWD and the ACC, e.g., the Prydz Bay and Ross Sea gyres (referencias). This region is a summer feeding ground for several large whale species as well as for other marine predators (Referencias. Lucho y Isida pueden ayudar).

Beyond their distinct oceanographic profiles, these Subantarctic and Antarctic regions share a critical ecological role as primary foraging grounds for different whale populations and other marine predator species (e.g., Acevedo et al. 2007, 2017; Constantine et al., 2014; et al., 2022). Consequently, characterizing the microbial baselines of these seawater regions not only advances Southern Ocean biogeography but also provides essential ecological insights into high-latitude marine ecosystems that are crucial for the energetic maintenance of global marine megafauna.

Here, we use high-throughput sequencing of 16S rDNA to characterize marine microbial communities across distinct seawater masses in two regions of the Southern Hemisphere used by several whale species as feeding areas: an open-sea Antarctic region and an inland Subantarctic region. Concurrently, marine oceanographic data were measured or obtained by satellite measurements to assess the relationship between environmental gradients and microbial diversity. We hypothesized that differences in oceanographic conditions between the Subantarctic and Antarctic regions would drive substantial differences in the ecological features of their respective marine microbial communities.

## Materials and Methods

### Study area

The target of this study was seawater masses of two Southern Hemisphere regions, which are important as feeding grounds for several marine predators including large whales: an Antarctic open sea region involving two localities (Eastern Indian Ocean [70°00’ - 129°36’E, south of 60° S] and Central South Pacific Ocean [133°20’ - 143°54’W, south of 60° S]) and a Subantarctic region in the tip of South America involving two localities (Magellan Strait [53°30’S; 72°15’W] and the Chilean sector of Beagle Channel/Cape Horn [54°54’S; 68°25’W]) (Figure 1).

**Figure 1.**
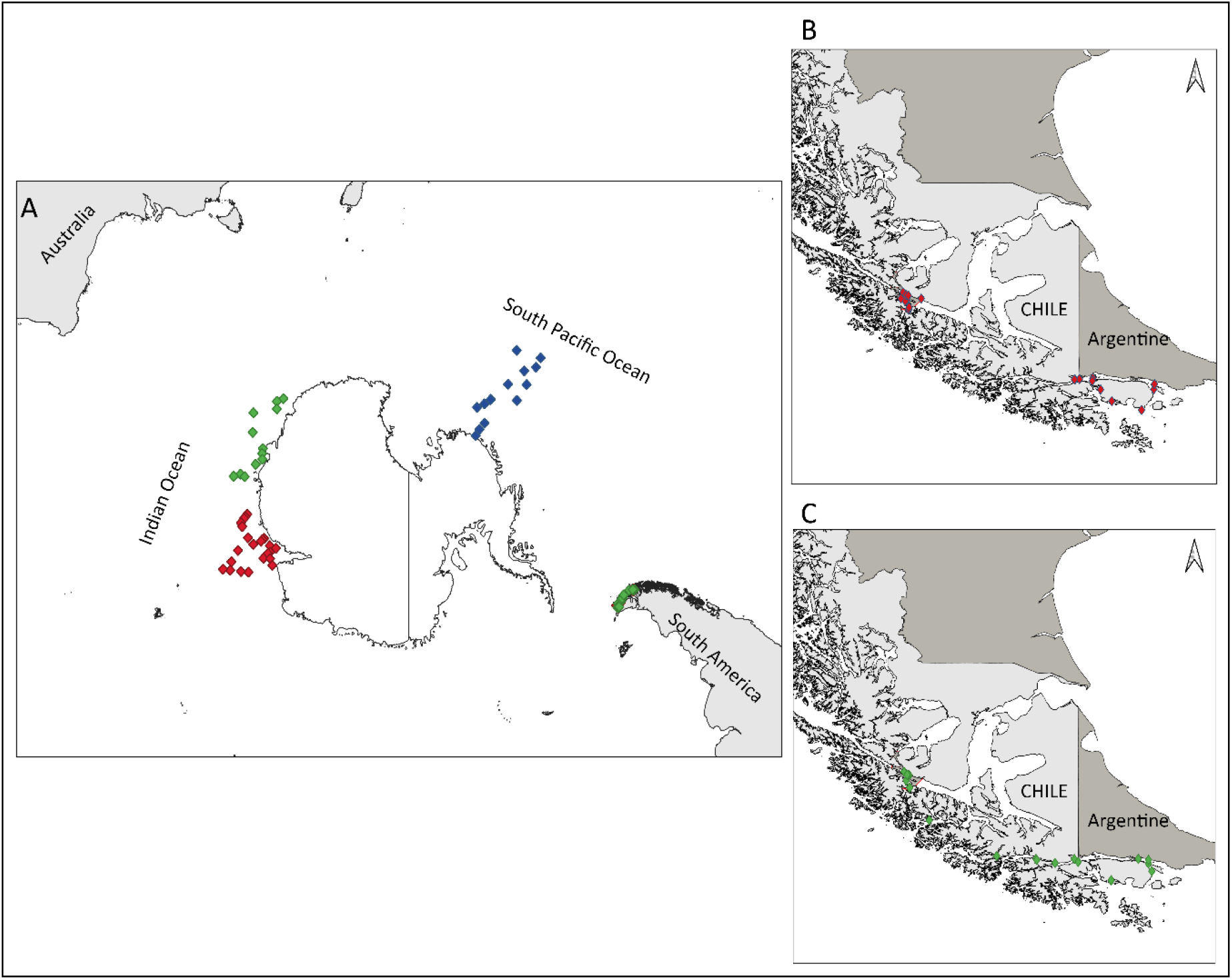
Distribution of the seawater samples across years in the Antarctic Indian and Central South Pacific Oceans (A), and the Subantarctic region of Magellan Strait and the Chilean sector of Beagle Channel (B and C, respectively). Blue diamonds are samples collected in 2023, red diamonds in 2024, and green diamonds in 2025, respectively.

### Sample Collection and Processing

Overall, 98 surface seawater samples were analyzed. For the Antarctic region, 35 samples from the eastern Indian Ocean locality collected during January and February of austral summer seasons 2023/2024 and 2024/2025, and 14 samples from the central South Pacific locality collected during January and February of the 2022/2023 austral summer season, by the surveys of the Japanese Antarctic Abundance and Stock structure (JASS-A) program were used. For the Subantarctic region, 27 samples from the Magellan Strait collected between January and May of the 2024 and 2025, and 22 other samples from the Chilean sector of the Beagle Channel locality in April 2024 and February 2025 (summer/autumn seasons) by CEQUA surveys, were used.

Surface seawater samples were collected using bottles pre-washed with detergent and subsequently rinsed with seawater by submersion two or three times before taking the final samples. In the Antarctic region, 4L of seawater was immediately filtered onboard the vessel using an aspirator through a polytetrafluoroethylene (PTFE) membrane filter (Millipore, Inc) with a pore size of 0.45 μm and a 90 mm diameter. In the Subantarctic region, 3L of seawater was immediately filtered onboard the vessels using a vacuum pump through a polycarbonate membrane filter (Advantec, MFS Inc.) with a pore size of 0.2 μm and a 47 mm diameter, allocating 1 L of seawater per individual filter. To minimize potential background contamination, personnel strictly wore gloves and face masks throughout all processing stages. All filters were immediately stored in liquid nitrogen and subsequently transferred to an ultra-low temperature freezer at -80°C at the respective laboratories of the Institute of Cetacean Research (ICR) and CEQUA until processing.

Concurrently, date, geographic position, and environmental variables were recorded at each sampling locality. In the Subantarctic region, sea surface temperature (SST, °C), chlorophyll concentration (mg m^-3^), salinity (psu), and dissolved oxygen (mg L^-1^) were recorded at depths ranging from 2 to 3 m using a WIMO NKE multiparameter probe. In the Antarctic region, SST was recorded using a sea water temperature sensor (Nippon Electric Instrument, Inc., Model N66M) mounted on the ship’s hull, at depths of 2 to 3 m. The other oceanographic variables were obtained from daily satellite-derived products available through the Copernicus Marine Data Viewer (https://data.marine.copernicus.eu/viewer/). Specifically, chlorophyll-a concentration was extracted from the Global Ocean Colour product (BGC_4L_MY_009_104) with a spatial resolution of 4 km, whereas sea surface salinity and dissolved oxygen were obtained from the Global Ocean Physics Analysis (PHY_001_024) and Forecast and Global Ocean Biogeochemistry Analysis and Forecast (BGC_001_028) products with spatial resolutions of 0.083° and 0.25°, respectively. Although these satellite-derived products differ in spatial resolution, each variable was extracted from the grid cell containing the geographic coordinates of each seawater sampling for the corresponding sampling date, providing a standardized representation of the oceanographic conditions at the time of sampling. Previous studies have shown that Copernicus satellite and model products provide reliable estimates of sea surface chlorophyll-a, salinity, and dissolved oxygen, supporting their use in comparative ecological analyses (Stramska et al. 2021, Lamouroux et al. 2025).

Environmental variables from the two regions were analyzed separately. Given the heterogeneous variance and the non-normal distribution of the data, the Kruskal-Wallis test was used to analyze each environmental variable across each region’s localities. Upon significant results, the Wilcoxon pairwise test was conducted with Holm p-value correction for multiple comparisons.

### Laboratory work

#### DNA extraction

For the filter samples of the localities in the Antarctic region, DNA extraction was carried out at the genetic laboratory of the ICR, Japan. Total genomic DNA was extracted using a sample of 4L of seawater per filter. Lysis Solution F (Nippon Gene Co., Ltd., Toyama, Japan) was added to each filter, and the sample was disrupted using a Shake Master Neo (BMS Co., Ltd., Shinjuku, Tokyo, Japan) at 1500 rpm for 2 min. The disrupted samples were incubated at 65 °C for 10 min, followed by centrifugation at 12,000 g for 1 min-2 min, after which the supernatant was collected. DNA was then purified from the collected supernatant using a Lab-Aid 824s DNA Extraction Kit (Xiamen Zeesan Biotech Co., Ltd., Xiamen, China). The concentration of each DNA solution was measured using a Synergy LX microplate reader (Agilent Technologies, Inc., Santa Clara, CA, USA) with the QuantiFluor dsDNA System (Promega Corporation, Madison, WI, USA). Sequencing libraries were prepared using a two-step tailed PCR method. The concentration of each prepared library was measured using a Synergy H1 microplate reader (Agilent Technologies) with the QuantiFluor dsDNA System, and library quality was assessed using a Fragment Analyzer with the dsDNA 915 Reagent Kit (Agilent Technologies).

For the filter samples of the localities in the Subantarctic region, DNA extraction was carried out at the Genomic Laboratory of CEQUA in Punta Arenas, Chile. Total genomic DNA was extracted from an average sample of 1L of seawater per filter, using the Fast DNA Spin kit for soil (MP Biomedical, Santa Ana, Ca, USA) following a modified extraction protocol. Quantity and quality controls of DNA were determined using a Qubit 2.0 fluorometer (Life Technologies) and a Nanodrop spectrophotometer (Thermo Scientific), while DNA integrity was evaluated with gel electrophoresis. Extractions that failed initial quality control were re-extracted, and those showing degradation after extraction were size-selected for high molecular weight DNA using magnetic beads (Agencourt AMPure XP). To isolate microorganisms, 978 µl of Sodium Phosphate Buffer (SPB) was added to the samples and incubated in a MultiTherm heat shake (Benchmark Scientific, Inc., Edison, N.J., USA) at 1500 rpm and 56°C for 2 hours. For the homogenizing process, a FastPrep-24 5G bead beating device (MP Biomedicals, Irvine California, USA) was used at 4 m/s, with a time reduction of 30 s for two cycles, and an added 1 min ice bath between cycles, followed by centrifugation at 14,000 g for 15 min. For the protein precipitate step, samples were incubated at -20°C for 5 min before 10 min of centrifugation. To avoid residual washing solution, tubes were air-dried in a PCR cabinet for 30 min. To improve DNA concentration, the binding matrix was resuspended in 60 µl of Ambion Nuclease-Free Water (Invitrogen, Thermo Fisher Scientific) followed by a 5 min incubation step in the MultiTherm heat shake (Benchmark Scientific, Inc., Edison, N.J., USA) at 56°C.

To assess microbial contamination of kit reagents and the reliability of sequence processing and taxonomic assignments, distilled water blanks were processed for DNA extraction. In addition, two mock microbial communities of known species of Bacillus were isolated and sequenced along the seawater samples.

All DNA samples, including DNA blank controls and the two mock communities, were sent to Macrogen (Korea and Japan) for amplification of the hypervariable region V3–V4 of the 16S gene using the universal primers 341F-805R, library preparation, and paired-end (2 × 300 bp) sequencing using Illumina MiSeq.

### Bioinformatic analysis

The software DADA2 v1.18.0 was used in R to describe Amplicon Sequence Variants (ASVs) from raw reads (Callahan et al., 2016; R Core Team, 2022). Primer sequences from forward and reverse raw reads were removed in R. To obtain ASVs, ambiguous bases were not allowed, and a maximum of 1 (for forward) and 2 (for reverse) errors were allowed. Error rates for learning, dereplication, denoising, and merging of paired-end reads were conducted with default parameters. Taxonomic classification was done with the naïve Bayesian classifier (Wang et al., 2007), using the Silva 138.2 training set up to species database as reference (Quast et al., 2013). Sequences classified as Chloroplasts and Mitochondria, and those with low prevalence (i.e., only present in one sample) were discarded. The denoised sequences, the taxonomic classification, and the corresponding metadata were merged into a phyloseq object (McMurdie and Holmes, 2013)

The complete set of raw ASVs was normalized to a fixed sampling depth (32, 000), since it is the best procedure to prevent type I error when conducting diversity analyses (McKnight et al., 2019). According to rarefaction curves, at 32,000 sequences, all conditions reached the plateau (Supp. Fig. 1). However, upon conducting sequencing depth normalization, we lost one sample that had poor sampling depth (>11, 000 sequences).

### Community analysis

The 10 most abundant bacterial genera were aggregated and visualized by locality and year using the *microViz* R package (Barnett et al., 2021). Also, from the sequences that were annotated at the species level, the most abundant features were explored, and the genera with the highest number of species detected. To determine the bacterial core among regions (90% minimum prevalence and 0.1% minimum relative abundance), the core function of the microbiome package was used (Lahti and Shetty 2017).

Alpha diversity was measured with the Shannon index and observed richness of ASVs calculated with the phyloseq package. Differences in alpha diversity among regions and years were assessed using Kruskal–Wallis tests. Upon significant results, the post hoc Wilcoxon rank sum test, using Holm p-value adjustment for multiple comparisons, was used to address specific differences among conditions. To determine if there were correlations between Shannon values and environmental factors, we conducted Spearman correlations across localities within each region. We kept significant correlations with p values > 0.05, but also show clear tendencies that do not fall under this significance criterion (p values between 0.05 – 0.06).

To determine significant variation in bacterial composition across regions and localities, the Bray-Curtis dissimilarity index was used, followed by the permutational analysis of variance (PERMANOVA) (Anderson, 2017). When significant differences were detected, post hoc paired PERMANOVA tests with p-value adjustment for multiple comparisons were conducted to detect specific differences among conditions. Distance to the centroid for each factor was extracted using the beta.disper function and statistically tested with the permutest function from the vegan package (Oksanen et al., 2022), which provides insight into the adequacy of the PERMANOVA results (Anderson and Walsh, 2013). As a complementary approach, BC similarity values across conditions were calculated and analyzed using a Bayesian approach with the brms R package (Bürkner, 2017a). Since similarity values (response variable) are bound between 0 and 1, a beta regression was used. Uninformative priors were used in all models. Models estimated in the brms package use a Markov Chain Monte Carlo sampler, implemented with Rstan (Stan Development Team, 2023), to estimate posterior distributions (Bürkner, 2017b). Each model was implemented with four parallel chains, each chain with 1000 warm-up samples embedded in 4000 iterations. The convergence of each was diagnosed, ensuring that Rhat values were equal to 1 and bulk effective samples were equal to or greater than 10% of total posterior draws. Also, posterior estimated parameters (mean and standard deviation) were checked to ensure their resemblance to the observed values (Gabry et al. 2019). The estimated mean and 95% credibility intervals were plotted within each region across years.

To obtain enriched taxa across region/localities and years and to find taxa with significant relationships with oceanographic variables, we used the software microbiome multivariable associations with linear models (MaAsLin3, Nickols et al., 2024). MaAsLin3 uses a GLM to model taxa abundance with selected categorical and continuous factors; it log-transforms taxa abundance and uses total sum scaling normalization. Statistical significance from individual associations was corrected with the false discovery rate. Hence, we use the non-rarefied dataset and two models to find patterns at different ecological scales. To find enriched taxa between the Antarctic and Subantarctic regions, the model was: taxa ∼ Ecological_region + Reads, while to find enriched taxa across localities within each region and taxa with significant correlations with oceanographic variables, the model was: taxa ∼ Ecological_region + SST + Chlorophyll + Salinity + Oxygen + Reads. We applied these models to two sets of datasets, one with the most precise taxonomic classification (ASV-level), and one with taxonomy agglomerated at the genus level, to determine if patterns were idiosyncratic within genus or were consistent at each taxonomic category.

## Results

### Environmental factors across regions and localities

Significant variability in environmental factors showed distinct patterns across regions (Supp. Fig. 1). In the Antarctic region, SST and Salinity were similar across localities (SST: KW = 3.510, p = 0.172; Salinity: KW = 2.179, p = 0.336). In contrast, SST and Salinity had significant variation across localities in the Subantarctic region (SST: KW = 12.59, p = 0.005; Salinity: KW = 14.213, p = 0.002). In the SST, post hoc tests revealed that the conditions with the greatest differences were the Beagle Channel during 2024 and the Magellan Strait during 2025 (p.adj = 0.005). Also, a significant tendency was observed in the Beagle Channel across years (p.adj = 0.055). In Salinity, post hoc tests revealed that the conditions with the greatest differences were between the Beagle Channel during 2024 and the Magellan Strait during 2024 (p.adj = 0.006) and during 2025 (p.adj = 0.015). Chlorophyll concentration significantly differed across localities in the Antarctic region (KW = 15.172, p > 0.001), specifically between the Central South Pacific and the Indian Ocean during 2024 (p.adj = 0.001) and 2025 (p.adj > 0.001). In contrast, chlorophyll concentration was similar across localities in the Subantarctic region (KW = 2.462, p = 0.482). Finally, dissolved oxygen differed in the Antarctic (KW = 16.784, p > 0.001) and Subantarctic regions (KW = 9.025, p = 0.028). In the Antarctic region, the main differences were between the Central South Pacific locality and the Indian Ocean during 2024 (p.adj > 0.001). In turn, in the Subantarctic region, the main differences were during 2024 between the Beagle Channel and the Magellan Strait (p.adj = 0.019).

**Table 1.** Environmental factors across regions, localities, and years. SST stands for Sea Surface Temperature.

| Antarctic region |  |  |  |  |  |
| --- | --- | --- | --- | --- | --- |
| Locality | Year | SST (°C) | Chlorophyll (mg/m<sup>3</sup>) | Salinity (psu) | Dissolved Oxygen (mg/L) |
| Central South Pacific | 2023 | -0.02 ± 0.3 | 0.4 ± 0.2 | 33.4 ± 0.3 | 11.7 ± 0.2 |
| Eastern Indian Ocean | 2024 | 0.5 ± 0.8 | 0.6 ± 0.6 | 33.4 ± 0.4 | 11.4 ± 0.1 |
|  | 2025 | 0.6 ± 1.2 | 0.1 ± 0.1 | 33.5 ± 0.5 | 11.6 ± 0.2 |
| Subantarctic region |  |  |  |  |  |
| Beagle Channel | 2024 | 8.1 ± 0.3 | 1.2 ± 0.4 | 29.9 ± 0.7 | 8.8 ± 0.4 |
|  | 2025 | 8.9 ± 0.7 | 1.6 ± 0.6 | 28.2 ± 1.9 | 9.8 ± 1 |
| Magellan Strait | 2024 | 8.5 ± 0.4 | 1.6 ± 1.1 | 29.2 ± 0.3 | 9.8 ± 1 |
|  | 2025 | 8.9 ± 0.3 | 1.2 ± 0.3 | 28.9 ± 0.4 | 9.6 ± 0.9 |

### Taxonomic patterns

Overall, the rarefied dataset comprised 3,072,000 reads across 20 phyla, 36 classes, 89 orders, 186 families, and 302 genera. In total, 2,122 taxa were identified, of which 126 were annotated to the species level (5.9% of the total). The most abundant genera across regions and years were *Clade Ia*, *Planktomarina*, *Polaribacter*, and *Yoonia*. *Clade Ia* was the most dominant genus in most of the localities and years. It was particularly abundant in the Antarctic region (eastern Indian Ocean locality) during 2025 and in the Subantarctic region (Beagle Channel locality) during 2025, where it reached relative abundances of 22.94% and 15.77%, respectively (Fig. 2). Other dominant genera in the Antarctic region included *Yoonia* (24.87%) (central South Pacific locality) during 2023 and *Polaribacter* (20.06%) (eastern Indian Ocean locality) during 2024. Besides these patterns, four genera comprising a strict spatiotemporal core (prevalent in 90% of samples) were detected: *Clade Ia*, *Amylibacter*, *NS5 marine group*, and *NS2b marine group*.

**Figure 2.**
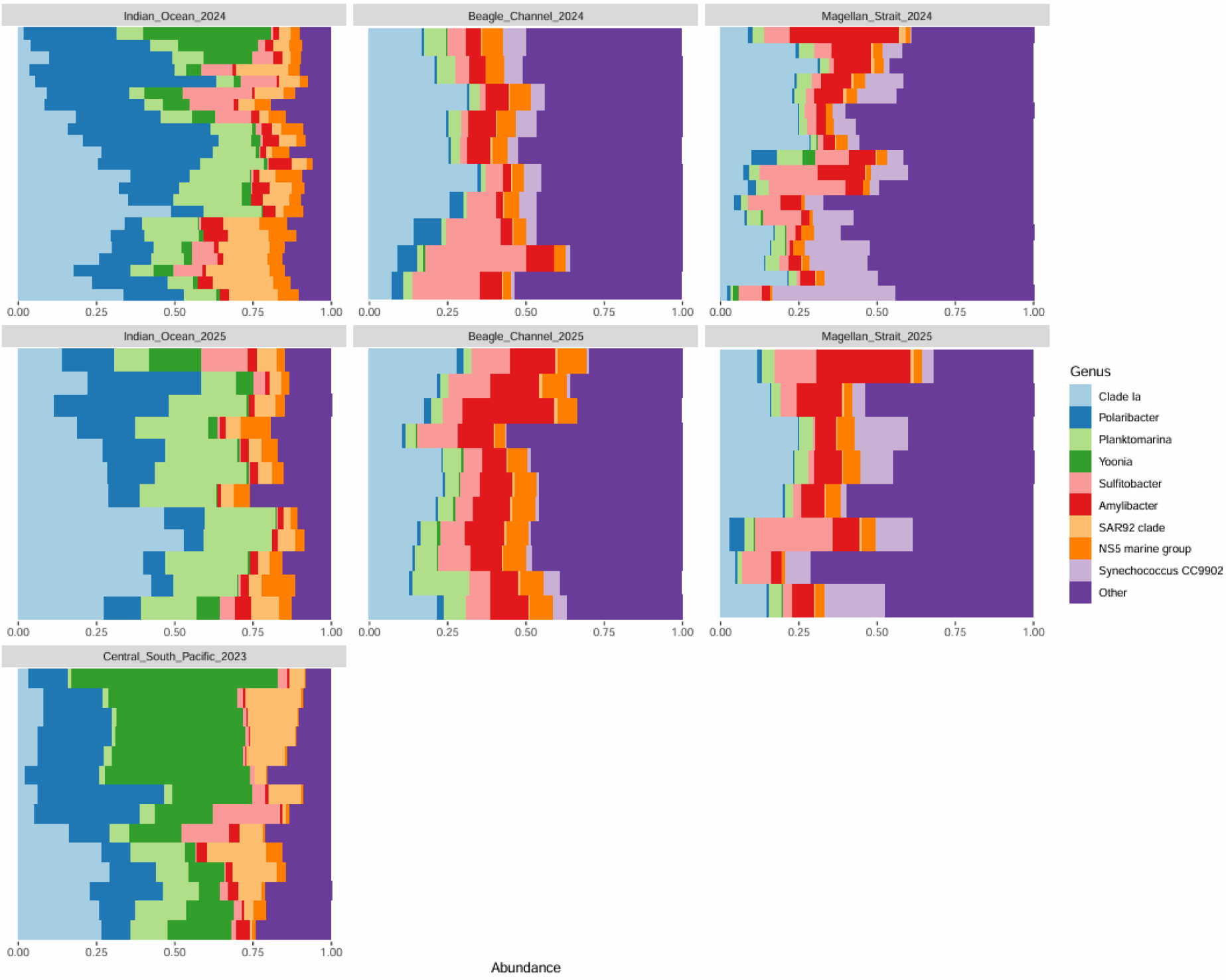
Interannual taxonomic patterns in the bacterial communities at the genus level across Antarctic and Subantarctic regions and localities.

In terms of species-specific taxa, different patterns were detected across regions and localities. In the Subantarctic region, during both years, the most abundant species was *Alteromonas stellipolaris*, followed by *Croceibacter atlanticus* and *Nocardioides salarius*. Yet, there were other relevant species in this region, such as *Idiomarina Ioihiensis*, *Alcanivorax venustensis*, and *Vibrio splendidus*. Conversely, from the ASVs that could be annotated at the species level in the Antarctic region, the most abundant was *Polaribacter irgensii*, while the other species in this region were present at low relative abundances (>0.1%). Globally, among the genus with more annotated species in both regions (Subantarctic and Antarctic regions), the most notable are *Psychrobacter* (six species: *P. nivimaris*, *P. marincola*, *P. urativorans*, *P. cibarus*, *P. pacificensis*, and *P. piscatori*), *Glaciecola* (four species: *G. pallidula*, *G. punicea*, *G. siphonariae*, *G. nitratireducens*), and *Colwellia* (three species: *C. rossensis*, *C. beringensis*, *C. psychrerythraea*). A complete list of the annotated species is provided as supplementary material (Supp. Table 1).

### Dynamics in marine microbial alpha and beta diversity

Significant differences in both the Shannon index and ASV richness were detected among regions and localities, where alpha diversity was higher in the Subantarctic localities (Fig. 3a, Shannon: KW = 67.553, p > 0.0001; Fig. 3b, ASV richness: KW = 75.252, p > 0.0001). Post hoc pairwise comparisons revealed that Shannon values differ significantly between Subantarctic localities, where the Beagle Channel locality displayed higher microbial alpha diversity than in the Magellan Strait locality in both years (Fig. 3a, Supp. Table 2). Also, Shannon values significantly differed between the Antarctic localities, where the eastern Indian (in any year) displayed higher microbial alpha diversity than in the central South Pacific of 2023 (Fig. 3a, Supp. Table 2). In contrast, Shannon index values were similar across years within localities in the Subantarctic region (e.g., Magellan Strait 2024, 2025) and Antarctic regions (Supp. Table 2). ASV richness patterns were similar to those found in Shannon values, where the Subantarctic locality of the Beagle Channel displayed the highest richness from all sites in both years, while the ASV richness from the Antarctic region remained similar despite spatiotemporal variability (Fig. 3b).

**Figure 3.**
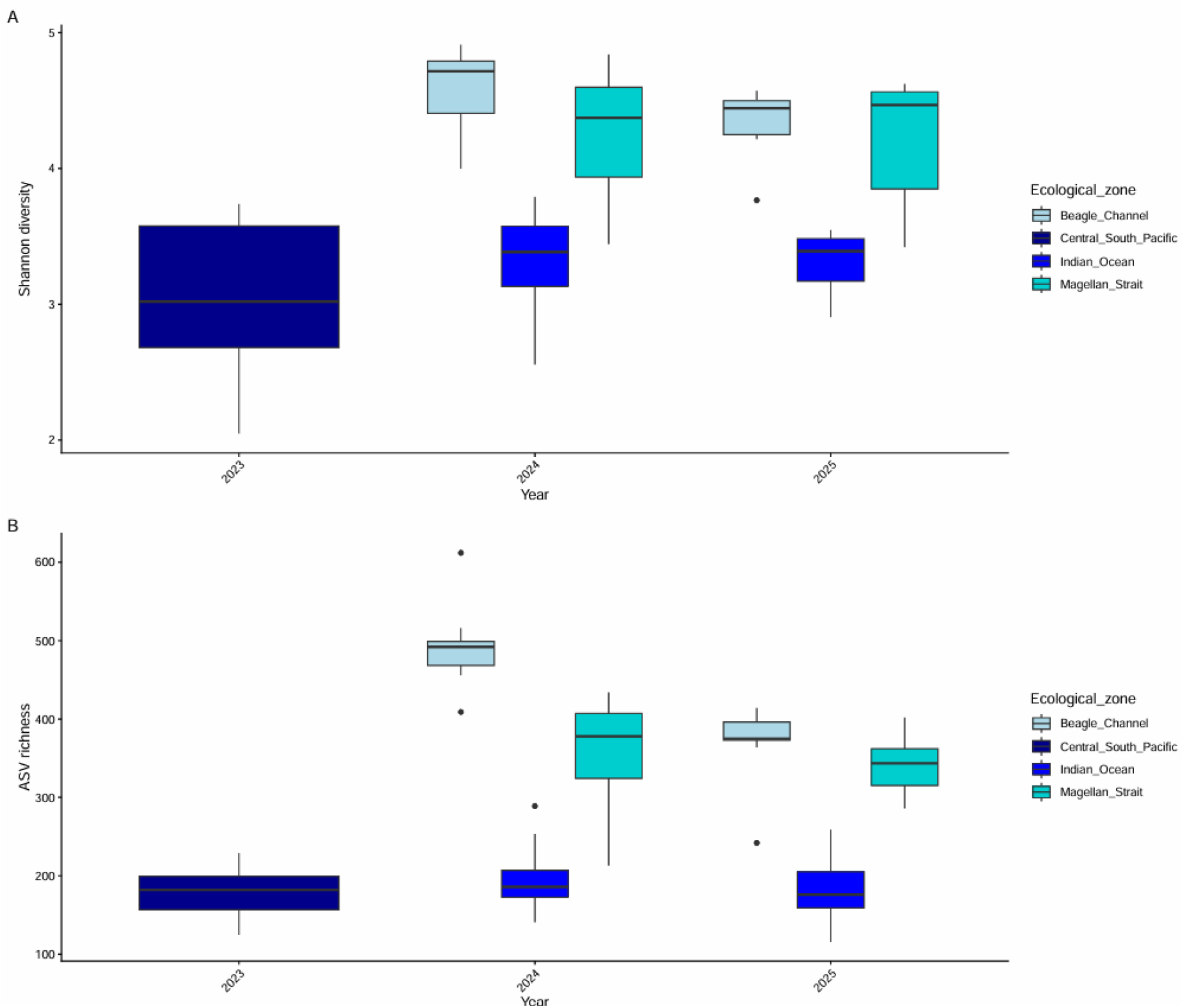
Spatiotemporal patterns in microbial alpha diversity across Antarctic and Subantarctic regions. A) Shannon diversity, B) ASV richness.

Global ordination analysis revealed strong compositional differences (almost 50% of variance explained in the first axis of the PCoA) between Subantarctic and Antarctic seawater microbial communities (Fig 4a, pseudo F = 84.9999, R^2^ = 47.48, p = 0.001). Variability in calculated community similarity followed these compositional differences, where microbial communities in the Subantarctic region (66.62%) were 4% more similar among them than those in the Antarctic region (62.47%) (Fig 4d). However, subtle differences in dispersion between Antarctic and Subantarctic samples raise caution in the adequacy of the PERMANOVA result (betadisper, F = 4.17, p = 0.04). Post hoc pairwise PERMANOVA tests revealed that in all pairwise comparisons, microbial communities had significant differences in their compositions (Supp. Table 3).

**Figure 4.**
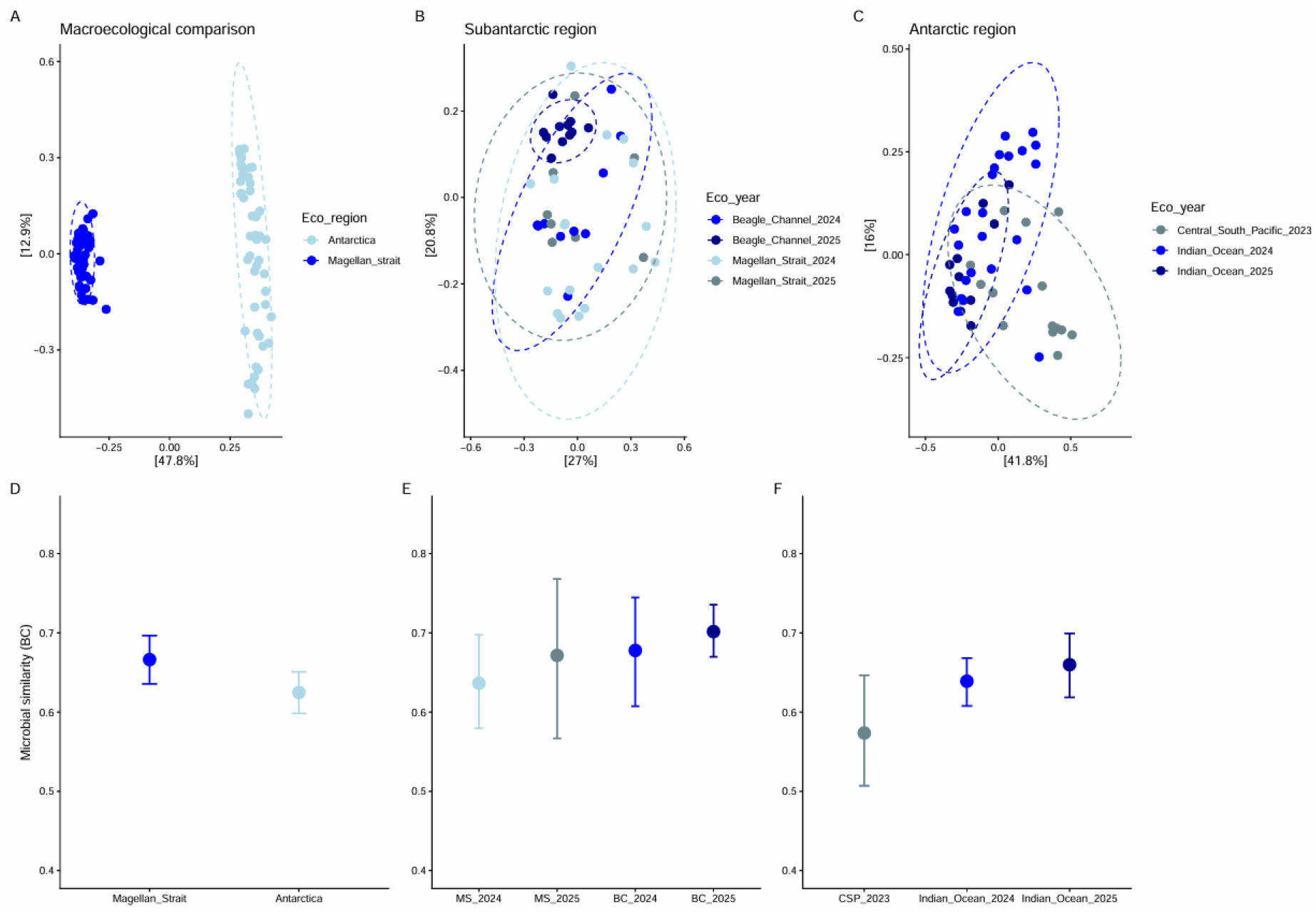
Spatiotemporal patterns in the seawater microbial composition and similarity in the Southern Ocean. Principal Coordinate Analysis using the Bray-Curtis dissimilarity index of marine microbial communities at various levels: A) Antarctic vs Subantarctic regions; B) Within the Subantarctic region: Magellan Strait and the Beagle Channel localities through 2024 and 2025; C) Within the Antarctic region, Central South Pacific (2023) and eastern Indian Ocean localities (2024 and 2025). Microbial similarity comparisons at the same spatiotemporal scale as the ecological ordinations D-F. In D – F, dots show estimated average similarity based on the Bray–Curtis distance, and lines represent 95% credibility intervals

Within regions, the microbial compositional analysis revealed significant spatiotemporal differences. In the Subantarctic region, seawater microbial communities had distinct compositional differences across localities and years (Fig. 4b, pseudo F = 3.3939, R^2^ = 19.14, p = 0.001). These differences reflect genuine microbial compositional differences, rather than differences in dispersion (betadisper, F = 2.517, p = 0.08). However, further PERMANOVA pairwise comparisons revealed that the seawater of the Magellan Strait locality had a more stable community among years (F = 1.230, R^2^ = 0.04, p = 0.27), which contrasts with the pattern observed in the Beagle Channel subantarctic locality, where there were significant compositional differences across years (Pseudo F = 5.208, R^2^ = 0.215, p = 0.002). Interannual comparisons between subantarctic localities, that is, Beagle Channel vs Magellan Strait, consistently showed significant differences (Supp. Table 3). Likewise, community similarity exhibited interannual variability (Fig. 4e). In the Magellan Strait locality, the seawater community similarity increased by approximately 4%, from 63.64% in 2024 to 67.14% in 2025. In the Beagle Channel locality, interannual seawater community similarity variation was less pronounced, increasing by about 3% from 67.77% in 2024 to 70.15% in 2025.

A similar geographic pattern, although more intense since more variance was explained by geographic variability (24.96% vs 19.14% in the Subantarctic region), was detected in the Antarctic region, where seawater microbial communities harbored significantly different compositions across localities and years (Fig. 4c, pseudo F = 7.65, R^2^ = 24.96%, p = 0.001). These differences reflect genuine microbial compositional differences, rather than differences in dispersion (betadisper, F = 3.24, p = 0.056). Further PERMANOVA pairwise comparisons revealed consistent differences among years within the eastern Indian Ocean during 2024 in relation to 2025, and across localities (central South Pacific vs eastern Indian Ocean) (Supp. Table XX). These compositional differences were accompanied by stronger differences in community similarity, particularly between the central South Pacific 2023 and the eastern Indian Ocean 2025, where community similarity increased by > 8% from 57.36% in 2023 to 65.97% in 2025. In turn, in the eastern Indian Ocean locality, community similarity increased by> 2%, from 63.89% in 2024 to 65.97% in 2025.

### Effect of environmental factors on microbial alpha diversity

Environmental factors had a variable relationship with marine microbial diversity across regions (Fig. 5). In the Subantarctic region, particularly in the Magellan Strait locality during 2024, Shannon diversity had a negative correlation with SST (Fig. 5a, Rho: -0.653, p = 0.004), and a positive correlation with salinity (Fig. 5b, Rho: 0.591, p = 0.015). In contrast, in the Beagle Channel locality, there was no detectable relationship between any environmental factor and microbial diversity. In turn, in the Antarctic region, particularly in the eastern Indian Ocean locality during 2024, Shannon diversity had a negative tendency with dissolved oxygen (Fig. 5c, Rho: -0.407, p = 0.053). Also, in the eastern Indian Ocean locality, Shannon diversity had a positive correlation with salinity in 2024 (Fig. 5d, Rho: 0.606, p = 0.002) and a negative tendency with salinity in 2025 (Fig. 5d, Rho: -0.559, p = 0.058), respectively. In the central South Pacific locality, there was no detectable relationship between any environmental variable and microbial diversity.

**Figure 5.**
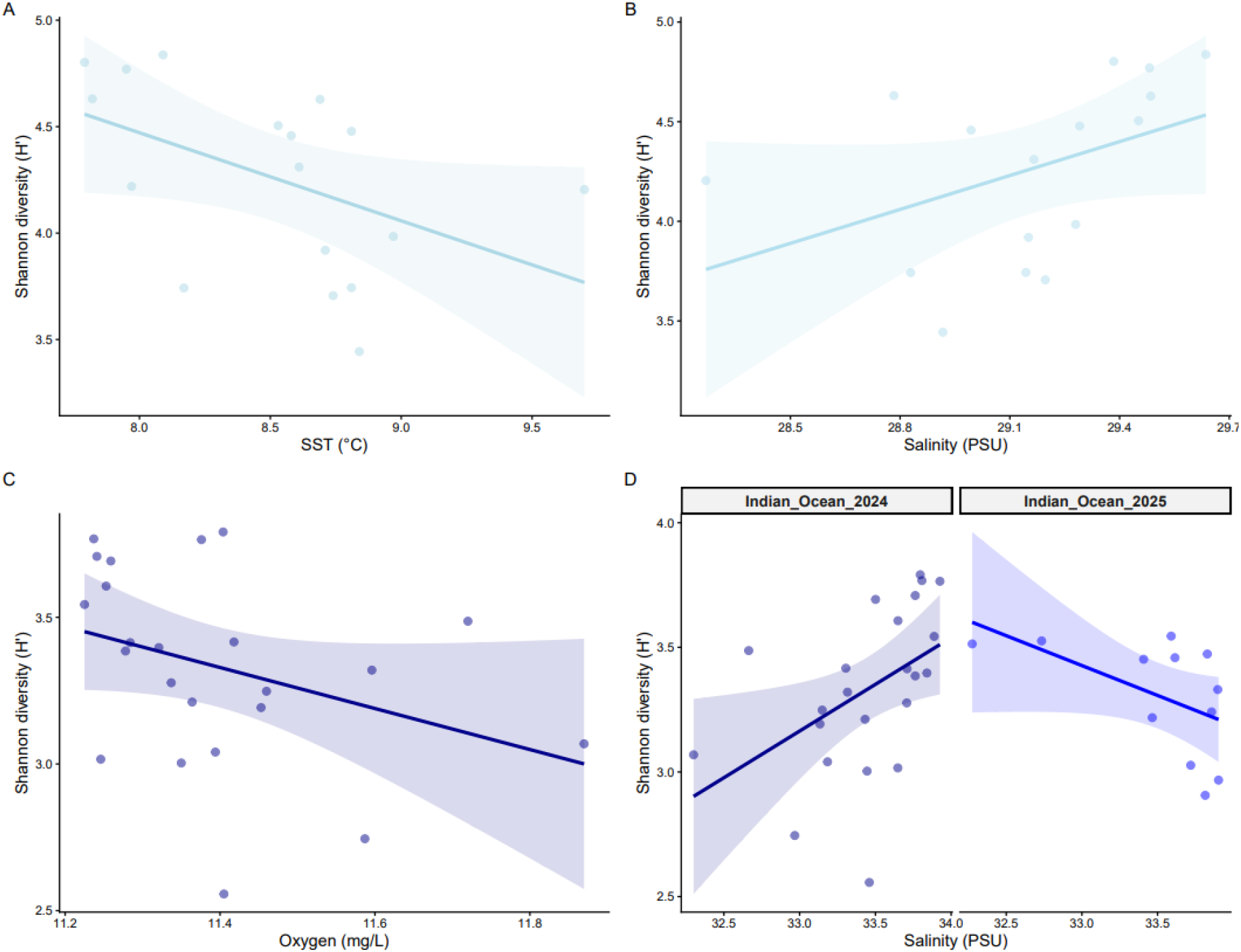
Relationship between microbial alpha diversity (measured with the Shannon index) and environmental factors. Correlations between microbial alpha diversity and sea surface temperature (SST) (A) and Salinity (B) in the Subantarctic region, particularly in the Magellan Strait locality during 2024. Correlations between microbial alpha diversity and dissolved oxygen (C) and Salinity (D) in the Antarctic region, particularly in the eastern Indian Ocean locality during 2024 and 2024 - 2025, respectively.

### Microbial signatures across regions in the Southern Ocean

Differential abundance analyses at the genus level between the Subantarctic and Antarctic regions revealed that *Candidatus Aquiluna*, *Psychromonas*, *Roseibacillus*, *CLP500-3*, and *Rubripirellula* were more abundant in the Subantarctic region. In contrast, *SAR92 Clade*, *Yoonia*, *Polaribacter*, *Octadecabacter*, and *Loktanella* were more abundant in the Antarctic region (Supp. Fig. 1).

Within-region differential abundance tests further clarify these differences. Within the Subantarctic region (Fig. 6), the most abundant taxa in the Beagle Channel locality were the bacterial families *PS1 clade* and *Nitrincolaceae*, and the bacterial genera: *IS-44* and *Luminiphilus*. In contrast, the most abundant taxa in the Magellan Strait locality were *Synechococcus CC9902* and *Pseudoalteromonas*. In the taxonomically agglomerated dataset, SST showed a positive relationship with the abundance of the families Pirellulaceae and Arenicellaceae, whereas Salinity showed a negative relationship with the family *S25-593*. ASV-level dataset further enriched the effect of environmental factors on microbial abundance (Supp. Fig. 3). Salinity was the oceanographic variable that affected the highest number of microbial taxa in the Subantarctic region; it had a positive relationship with *Clade II* (ASV 418) and *Clade Ia* (ASV 305), while it had a negative relationship with *Pseudohongiella* (ASV 47), *NS4 marine group* (ASV 17), and *Planktomarina* (ASV 5). In turn, SST had a positive relationship with *SAR 116 clade* (ASV 217) and a negative relationship with *Ascidiaceihabitans* (ASV 225).

**Figure 6.**
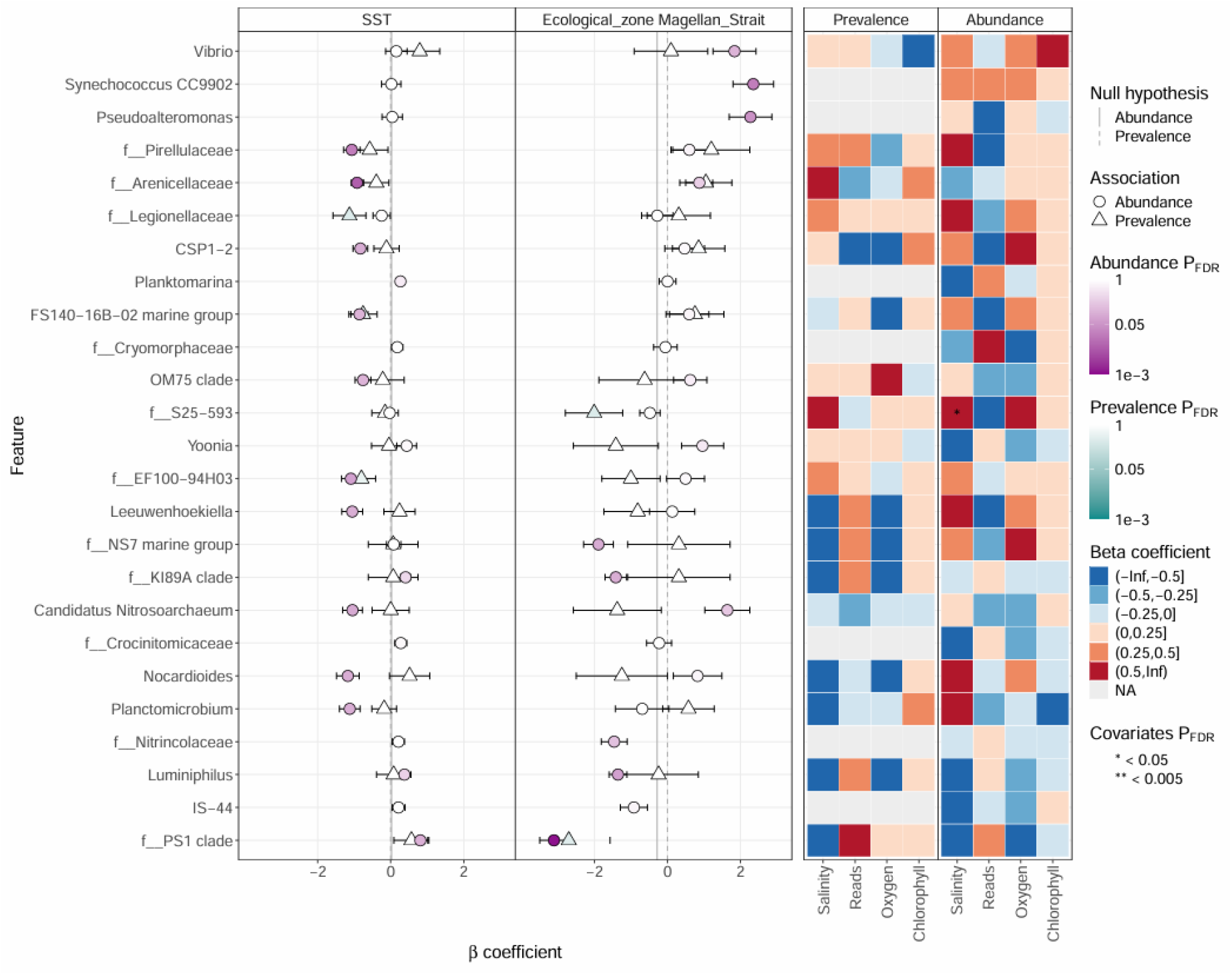
Patterns of differentially abundant taxa within the Subantarctic region. Results of MaASlin3 from enriched taxa as a function of Ecological zone, Salinity, SST, dissolved oxygen, and Chlorophyll concentration. Heatmaps reflect positive (red) and negative (blue) correlations. One asterisk indicates significance below 0.5. Taxa whose coefficient is negative are enriched either in the Beagle Channel locality or negatively associated with salinity, while those whose coefficient is positive are enriched either in the Magellan Strait locality or positively associated with salinity.

Within the Antarctic region (Fig. 7), the most abundant bacterial genera in the Central South Pacific locality were *Fluviicola*, *Yoonia*, *Octadecabacter*, and the *OM60(NOR05) clade*. In the eastern Indian Ocean locality, the most abundant genus was the *NS5 marine group*. Chlorophyll concentration was the variable that affected the highest number of bacterial genera; it had a positive relationship with *Psychoserpens* and *Sulfitobacter*, while a negative relationship with *Clade II, NS7 marine group, NS4 marine group*, and *SAR 86 Clade*. Conversely, SST had a negative relationship with *SUP 05 cluster* and the bacterial family *Nitricolaceae* (ASV 1292), while it had a positive relationship with *Ascidiaceihabitans*. These results were largely consistent with the ASV-level plots (Supp. Fig. 4).

**Figure 7.**
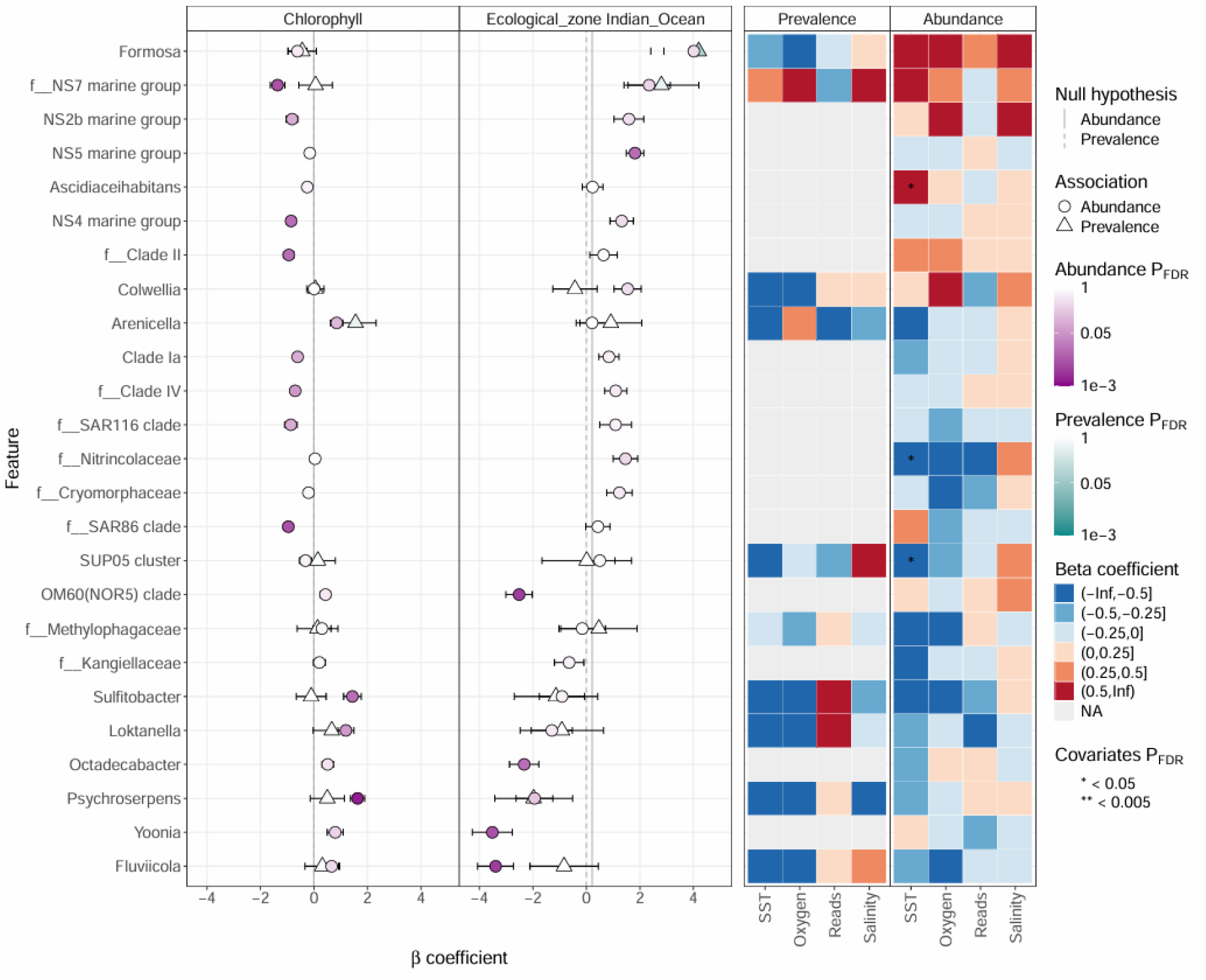
Patterns of differentially abundant taxa within the Antarctic region. Results of MaASlin3 from enriched taxa as a function of Ecological zone, Salinity, SST, dissolved oxygen, and Chlorophyll concentration. Heatmaps reflect positive (red) and negative (blue) correlations. Taxa whose coefficient is positive are enriched either in the eastern Indian Ocean locality or positively associated with Chlorophyll concentration, while those whose coefficient is negative are enriched either in the Central South Pacific locality or negatively associated with Chlorophyll concentration.

## Discussion

In this study, we presented for the first time a comprehensive characterization of seawater microbial diversity with next-generation sequencing approaches in two Subantarctic localities and two open-sea Antarctic localities, both regions widely used as feeding grounds by large marine predators.

### Dominant taxa across sites, core bacteria, and bacterial species insights

Bacterial genera such as *Clade Ia*, *Yoonia, Polaribacter*, and *Planktomarina* were the most abundant across Antarctic and Subantarctic regions. From the subset of ASVs that was possible to annotate to the species level, the most abundant in the Antarctic region was *Polaribacter irgensii*, while those in the Subantarctic region were *Croceibacter atlanticus*, *Nocardioides salaries*, *Alteromonas stellipolaris* (Alteromonadaceae), *Idiomarina Ioihiensis*, *Alcanivorax venustensis*, and *Vibrio splendidus*. Also, genera like *Psychrobacter*, *Glaciecola*, and *Colwellia* were relevant in terms of species richness. However, we also found a stringent core (90% prevalence) across subantarctic and Antarctic seawater masses, consisting of *Clade Ia*, *Amylibacter*, and *NS5 marine group*. These taxa are common members of the Southern Ocean; for example, *Clade Ia* and *Planktomarina* are among the most abundant taxa in the Southern Ocean (Raes et al., 2018). Also, common Antarctic bacteria like *Polaribacter*, *Yoonia*, and *Sulfitobacter* are routinely detected in Antarctic seawaters despite interannual variation (Pavlovska et al., 2025). Adaptations to endure oligotrophic conditions might facilitate their persistence in cold waters (Liu et al., 2023) or be positively correlated with blooms of primary productivity, like *Polaribacter* and *Colwellia* (Landa et al., 2018; Cordone et al., 2022). Interestingly, taxa like *Nocardioides*, *Psychrobacter*, and *Vibrio* are common members of the surface microbiota of marine hosts despite ecological variability, such as body-site, environmental exposure, and seasonality (Ochoa-Sánchez et al., 2023b, 2024, 2025a, 2025b). Overall, our results expand the geographic distribution of common taxa in Antarctic seawater and provide plausible evidence of the possible source of common microbial taxa found in marine hosts.

### Extensive differences in microbial alpha diversity and community composition across ecosystems

Microbial diversity and composition in the seawater show distinct patterns at distinct scales. There were strong differences in alpha diversity (both Shannon and richness) and microbial composition between the Subantarctic and Antarctic regions. Furthermore, microbial composition was 4% more heterogeneous in the Antarctic region than in the Subantarctic region. In the Subantarctic region, microbial alpha diversity was stable across years and locations, except for ASV richness, which was higher in the Beagle Channel locality during 2024. However, microbial composition had partial stability, as it was similar during both years at the Magellan Strait locality, yet it differed across Subantarctic localities at both years and displayed significant interannual variability. Interestingly, in the Beagle Channel locality, microbial communities displayed between 3 – 4 % more similarity than those in the Magellan Strait locality. Conversely, in the Antarctic region, microbial communities displayed similar alpha diversity profiles despite spatiotemporal variability, yet microbial compositions were unique at each site, with differences in microbial similarity up to ∼8% in the central South Pacific locality and ∼6% across Antarctic localities. Broad-scale differences might be the result of distinct patterns of marine productivity, which are positively associated with microbial diversity in the Southern Ocean (Raes et al., 2018). In this sense, Antarctic open seawater might have higher stability in its diversity patterns due to its oligotrophy, yet distinct oceanographic features like current dynamics and eddies might produce spatiotemporal microbial structure, which likely decreases microbial compositional similarity (Venkatachalam et al., 2017; King et al., 2023; Feng et al., 2022).

In contrast, the Subantarctic region of South America displays seasonal input of freshwater from rivers and glacier melt, which causes seasonal peaks of marine productivity in the region (Pantoja et al., 2011). Likewise, the more connectedness within Subantarctic localities and the repeated seasonal dynamics might positively influence microbial compositional similarity (Danczak et al., 2018). Our results add more evidence to the current knowledge on marine microbial biogeography in the Southern Ocean (Liu et al., 2020; O’Brien et al., 2022; Liu et al., 2024), indicating strong compositional differences between Subantarctic and Antarctic biomes, and across ecosystems within each biome.

### The interplay between Oceanographic variables and marine microbial diversity

Oceanographic variables affected marine microbial communities in diverse ways across Antarctic and Subantarctic regions. In the Subantarctic region, particularly in the Magellan Strait during 2024, microbial alpha diversity had a negative correlation with SST, while a positive correlation with salinity. In the Antarctic region, particularly in the eastern Indian Ocean, microbial alpha diversity had a negative correlation with dissolved oxygen concentration in 2024, while with salinity, it either had a positive correlation in 2024 or a negative correlation in 2025. Also, chlorophyll-a concentration, SST, and salinity had correlations with several taxa across regions in either positive or negative senses. These patterns might point to the importance of contingency on the starting community composition.

Salinity and dissolved oxygen concentration are strong drivers of microbial compositional turnover in cold waters and in Antarctic ice (Signori et al. 2014; Tortensson et al., 2015). Dissolved oxygen might affect microbial diversity in two ways: higher levels favor the proliferation of aerobic microbes (Ye and Shi, 2020), while at the same time restraining the proliferation of strict anaerobes; whereas lower levels might restrict aerobic microbes, while favoring anaerobic microbes (Jørgensen, 2000). Salinity changes require active metabolic accumulation of solutes to keep cellular osmotic balance (Dawson et al., 2023). Hence, whether salinity increases or decreases diversity might be contingent on the metabolic repertoire and the composition of the starting community (Piquet et al., 2011; Hernando et al., 2020). Indeed, given that our results revealed compositional shifts in the eastern Indian Ocean between 2024 and 2025, it is plausible that this compositional reconfiguration introduces microbes with limited capacity to cope with an environment that changes the salinity – diversity pattern, positive in 2024, while negative in 2025. It is also important to highlight the relevance of the ecological strategies of the microbes from the starting community, since copiotrophs might be favored by increases in chlorophyll-a concentration, while oligotrophs might be affected (Horner-Devine et al., 2003). Overall, these results highlight the dynamic interplay between marine microbial communities and oceanographic factors in the Antarctic and Subantarctic regions. These patterns might further affect microbial diversity in these regions, since environmental variables are expected to change in future decades, such as SST, which is expected to increase by at least 1°C in future years in the Subantarctic region (Ochoa-Sánchez et al., 2025b).

## Conclusions

Our work expands previous evidence, indicating the presence of profound differences at various levels in the marine microbial communities across Antarctic and Subantarctic regions. *Clade Ia* is the most abundant genus across regions, localities, and years, while in terms of richness of species detected, *Psychrobacter* is the most important, with six species detected. In terms of alpha diversity, the Subantarctic region displayed communities with higher microbial diversity than those in the Antarctic region. Similarly, microbial composition showed the greatest differences across regions, while also showing important differences across localities within each region. Interestingly, microbial compositional similarity was higher in the Subantarctic region, suggesting that in the Antarctic region there is higher compositional turnover across regions and years. Finally, we find a dynamic interplay between some oceanographic factors and microbial diversity and taxon-specific abundance, highlighting the importance of contingency in the starting community composition. Our work provides valuable insight into microbial-based biogeography in unexplored localities in the Southern Ocean and how oceanographic variables affect marine microbial diversity and abundance.

## Acknowledgments

We thank researchers, captains, and crew members who participated in the JASS-A surveys in the Antarctic region for their contributions to seawater sampling in the eastern Indian Ocean and central North Pacific localities. We also thank T. Sugimoto (ICR) for his work on DNA extraction of the JASS-A seawater samples. Samples of seawater in the Magellan Strait and Beagle Channel in the Subantarctic region were collected by CEQUA researchers in the context of the Microbiome Project funded by ANID, Chilean Government (R20F0009).

## Author contributions

Manuel Ochoa-Sánchez, Jorge Acevedo, Eliana P. Acuña-Gomez, and Luis Pastene conceived and designed the study. Yoshihiro Fujise, Tatsuya Isoda, Pedro Valenzuela, and Claudio Moraga performed sampling and laboratory experiments. Manuel Ochoa-Sánchez carried out the bioinformatics and statistical analyses. Manuel Ochoa-Sánchez wrote the first draft of the manuscript. Manuel Ochoa-Sánchez, Jorge Acevedo, Aida I. Murillo-Herrera, and Luis Pastene discussed the results and edited the paper. All authors read and approved the final version of the manuscript.

## Data availability statement

Raw 16S rRNA sequence data generated in this study have been deposited in the freely and publicly available NCBI Sequence Read Archive under BioProject PRJNA1512548.

